# Oligodendrocytes show enriched expression of amyloid precursor protein and GABA B receptor isoform 1a

**DOI:** 10.1101/2025.11.25.690528

**Authors:** Samah Houmam, Dominika Siodlak, Meena Seshadri, Gideon B. Hallum, Nathan P. Pezant, David R. Stanford, Casandra Salinas-Salinas, Yvonne M. Thomason, Courtney G. Montgomery, Heather C. Rice

**Affiliations:** Aging & Metabolism Research Program, Oklahoma Medical Research Foundation, Oklahoma City, OK, 73104, USA; Department of Biochemistry & Physiology, University of Oklahoma Health Sciences Center, Oklahoma City, OK, 73104, USA; Center for Geroscience and Healthy Brain Aging, University of Oklahoma Health Sciences Center, Oklahoma City, OK 73104; Center for Biomedical Data Sciences, Oklahoma Medical Research Foundation, Oklahoma City, OK, 73104, USA; Genes & Human Disease Research Program, Oklahoma Medical Research Foundation, Oklahoma City, OK, 73104, USA; Neuroscience Program, University of Oklahoma Health Sciences Center, Oklahoma City, OK, 73104, USA

**Keywords:** Oligodendrocytes | Amyloid Precursor Protein | APP | GABA Receptor | GABA_B_R | isoform-specific expression | Alzheimer’s disease

## Abstract

Amyloid precursor protein (APP) is a type I transmembrane protein that undergoes proteolytic processing to generate amyloid-β, the main component of amyloid plaques found in brains with Alzheimer’s disease. The proteolytic processing of APP also generates soluble APP alpha (sAPPα) which can modulate synaptic transmission and neurite outgrowth through the γ-aminobutyric acid type B receptor (GABA_B_R). Whether GABA_B_R mediates functions of sAPPα in other neural cell types such as glia remains unknown. sAPPα binds the R1a subunit isoform of GABA_B_R1 which contains two sushi domains absent in R1b. It is unclear whether both GABA_B_R1 isoforms are expressed equally across brain cell types. We determined relative RNA levels of the GABA_B_R1a and 1b isoforms in oligodendrocytes, microglia, endothelial cells, astrocytes, and neurons in adult mice using two approaches. We developed a GABA_B_R1 isoform-specific RNAseq analysis workflow to probe a publicly available dataset. We also isolated five cell types from a single mouse brain and performed RT-qPCR. We show that the GABA_B_R1a and 1b isoforms are differentially expressed among cell types. GABA_B_R1a expression was highest in oligodendrocytes and GABA_B_R1b expression was highest in astrocytes, suggesting that sAPPα-mediated GABA_B_R signaling may be most prominent in oligodendrocytes. We also confirmed that APP is expressed in all five cell types and showed that APP RNA levels are highest in oligodendrocytes. Together, our findings uncover cell type-specific expression of GABA_B_R isoforms and highlight oligodendrocytes as a principal cell type for GABA_B_R1a-mediated APP signaling, providing a foundation for future mechanistic studies.

## Introduction

The amyloid precursor protein (APP) is a type I transmembrane protein expressed in the brain as well as other organs^1^. APP was discovered in 1987 during a search to uncover the source of the amyloid beta peptide (Aβ), the main component of the amyloid plaques associated with Alzheimer’s disease (AD)^2^. APP undergoes proteolytic processing to release not only Aβ but also other fragments which have important roles in the brain. APP can be cleaved by either α- or β- secretase followed by γ-secretase. α-secretase cleaves within the Aβ region and thus precludes Aβ generation and produces soluble APP α (sAPPα), P3, and the APP intracellular domain (AICD). β-secretase initiates the amyloidogenic pathway which generates sAPPβ, Aβ, and AICD. APP can also be cleaved by η-secretase prior to α or β cleavage within the sAPP region to generate additional fragments APPη-α or APPη-β^3^. Full-length APP and its fragments are involved in crucial brain functions^4^. They modulate neurodevelopmental processes such as axonal outgrowth, dendritic arborization, neurogenesis, and neuronal migration^4,5^. In mature neurons, full-length APP and sAPPα regulate synaptic structure, plasticity, and neurotransmission^3,6^. Though best known for its neuronal roles, APP is also implicated in glial and endothelial functions^7^. APP KO mice display hypomyelinated axons^8^, a blunted microglial and astrocytic inflammatory responses^9^, mitochondrial fragmentation in astrocytes^10^, and reduced production of endothelial nitric oxide species^11^. Despite these observations, the role of APP in glial cells remains largely undefined and understudied.

Identifying proteins that interact with APP and its fragments has been a key strategy to elucidate its physiological functions^4^. In 2019, full-length APP^12^ and its secreted ectodomain sAPPα^13^ were shown to interact with the γ-amino butyric acid (GABA) type B receptor (GABA_B_R). GABA_B_R is a class C G-protein coupled receptor (GPCR) and is an obligatory heterodimer^14^. The GABA_B_R1 subunit contains the endogenous agonist binding site, and GABA_B_R2 is intracellularly linked to G-proteins. The GABA_B_R1 subunit has various isoforms with R1a and R1b being the main two variants responsible for GABA_B_R diversity in humans^14^. The main difference between the two isoforms is the presence of two sushi domains on the N-terminus of R1a but not in R1b due to differential promoter usage^15^. These sushi domains are responsible for the preferential localization of the GABA_B_R1a subunit to axon terminals^16,17^. The intact (full-length APP)^12^ and shed (sAPPα) ectodomain^13^ both bind at the N-terminal sushi domain 1 of GABA_B_R1a^13^. The sAPP-GABA_B_R interaction was shown to reduce synaptic vesicle release consequently modulating hippocampal synaptic plasticity and neurotransmission^13^ as well as neurite outgrowth^18^. Moreover, deletion of APP impaired GABA_B_R-mediated presynaptic inhibition and axonal GABA_B_R levels^12^. The downstream mechanism mediating the effects of sAPPα-GABA_B_R signaling remains under investigation as studies suggest it may deviate from canonical GABA_B_R signaling^19^.

In the brain, both GABA_B_R1a and GABA_B_R1b are expressed in most neuronal subpopulations including glutamatergic, dopaminergic, and GABAergic neurons^20^. Neurons utilize canonical GABA_B_R signaling pathways to regulate key processes like synaptic transmission and excitatory/inhibitory balance^14^. GABA_B_Rs couple to Gα_i/o_ and Gβγ proteins to inhibit adenylyl cyclase and presynaptic Ca2+ channels to reduce presynaptic release probability and activate postsynaptic GIRK channels to reduce neuronal excitability^14,20,21^.The inhibitory functions of GABA_B_R have been extensively studied in neurons but are not as well characterized in glial cells. Oligodendrocytes express functional GABA_B_R1 coupled to adenylyl cyclase^22^. Furthermore, GABA_B_R activation using Baclofen, an agonist with no isoform selectivity, regulates oligodendrocyte precursor cell differentiation in vitro^23^ and can accelerate remyelination after induced primary demyelination in mouse spinal cord^24^. Studies in astrocytes and microglia show that these glial cells also express GABA_B_R1 and its activation using baclofen resulted in a decreased inflammatory response to lipopolysaccharide (LPS)^25–27^. A study using Baclofen in endothelial cells suggests that they also express functional GABA_B_Rs which can modulate their response to homocysteine toxicity^28^. Although some of these studies detected both isoforms of GABA_B_R1, it is still unclear whether glial cells equally express and utilize both the GABA_B_R1a and R1b subunit isoforms, or if there are any preferences towards a specific isoform, which would provide insight into whether GABA_B_R1a may mediate functions of APP in other cell types besides neurons.

In this study, we provide a comprehensive cell type-specific expression profile of APP across oligodendrocytes, microglia, endothelial cells, astrocytes, and neurons sorted from the same mouse brain^29^. We further assessed isoform-specific GABA_B_R RNA expression in these five cell types using RT-qPCR. This study provides a basis for elucidating the role of GABA_B_R-mediated APP signaling across distinct brain cell types.

## Methods

### Animals

All experiments involving animals were conducted according to the Oklahoma Medical Research Foundation ethical guidelines and approved by the Institutional Animal Care and Use Committee. This study used C57Bl6 (Jax strain #000664) male and female mice aged 4-5 months (19-21 weeks). Age-matched male GABA_B_R1a KO and APP KO (Jax strain # 004133) mice were used. GABA_B_R1a KO mice were generated by the VIB-KU Leuven Center for Brain & Disease Research Mouse Expertise Unit, with support from VIB Discovery Sciences. GABA_B_R1a KO mice were provided by Joris de Wit, VIB-KU Leuven Center for Brain & Disease Research, Leuven, Belgium.

### RNA-seq analysis of publicly available datasets

The publicly available dataset analyzed in this study was generated by Zhang et al. (2014)^30^ (brainrnaseq.org) and is available on GEO under the accession number: <u>GSE52564</u>. To probe expression of GABA_B_R1 (ensembl ID: ENSMUSG00000024462) and GABA_B_R2 (ensembl ID: ENSMUSG00000039809), sequence alignment map (.sam) files were generated using the STAR aligner (2.7.11b) to align reads to the *Mus musculus* GRCm39 reference genome. To probe expression of GABA_B_R1a and GABA_B_R1b isoforms, we edited the GRCm39 reference genome file to annotate the transcript regions unique to each isoform as Gabbr1 (1a) and Gabbr1 (1b) respectively. The start and end of the isoform-specific exon regions were determined based on the sequences of the transcript ID ENSMUST00000025338.16 (Gabbr1-201) for Gabbr1 (1a) and ENSMUST00000172792.8 (Gabbr1-203) for Gabbr1 (1b). After each alignment, .sam files were converted to binary alignment map files (.bam) using samtools (1.18). Read counts were then generated using the featureCounts command in the subread package (2.1.1) and normalized using DESeq2 (1.42.1) in R.

### Magnetic cell-sorting of whole mouse brain

The cell-sorted fractions were prepared as described in Houmam et al. (2025)^29^. Briefly, mice were anesthetized with isoflurane and euthanized by cervical dislocation and decapitation. The brain was dissected then rinsed in ice-cold D-PBS with calcium, magnesium, glucose, and pyruvate (Thermo Fisher Scientific 14287080). A cut was made to remove the cerebellum, and the remaining cerebrum (forebrain and midbrain) was cut into small cubes on a cold block. The tissue was then transferred to a gentleMACS C-tube containing 5mL of dissociation buffer (papain in EBSS (Worthington LK003150) supplemented with trehalose (MilliporeSigma T9531-10G), AP-5 (MilliporeSigma 165304-5MG), Kynurenic acid (MilliporeSigma K3375-1G), and DNAse (Worthington LK003150)). Actinomycin-D (MilliporeSigma A1410-2MG), anisomycin (MilliporeSigma A9789-5MG), and triptolide (MilliporeSigma T3652-1MG) were added to inhibit ex-vivo activation of glial cells^31^. After dissociation, debris was removed using a gradient spin as in the Worthington dissociation kit, and the solution was strained through a 70um filter to prepare a single-cell suspension in DPBS with 0.5% BSA (Fisher BP1600-100). This was used as input in subsequent RT-qPCR analyses. Myelin removal beads (Miltenyi 130-096-733) were used to separate the suspension into a myelin-positive fraction from which oligodendrocytes were isolated using anti-O4 beads (Miltenyi 130-096-670), and a myelin-negative fraction. From the negative fraction we sequentially isolated microglia using anti-CD11b beads (Miltenyi 130-126-725), endothelial cells using anti-CD31 beads (Miltenyi 130-097-418), astrocytes using anti-ACSA-2 beads (Miltenyi 130-097-678), and neurons using negative selection with biotin tagged beads (Miltenyi 130-115-389). The cell-sorted fractions were validated using RT-qPCR and flow cytometry for cell-specific markers^29^.

### RNA extraction and RT-qPCR

RNA was isolated using the Qiagen RNeasy Mini Kit (QIAGEN 74104) following the kit instructions, then quantified using a Nanodrop 1000 spectrophotometer. RNA quality was assessed using a 4150 Tapestation analyzer. RNA was converted to cDNA using the High-Capacity cDNA Reverse Transcription Kit (Applied Biosystems 4374966) in a BioRad T100 thermal cycler. RT-qPCR was performed in 384-well plates using Taqman primers and Gene Expression Master Mix (Applied Biosystems 4369016) following the manufacturer instructions. Commercially available taqman primers were used to probe for GABA_B_AR1(Mm00444578_m1) and GABA_B_R2 (Mm01352554_m1). Primers used to probe for GABA_B_R1a and GABA_B_R1b transcripts were designed using the <u>ENSMUST00000025338.16</u> and <u>ENSMUST00000172792.8</u> sequences respectively.

**Table 1:**
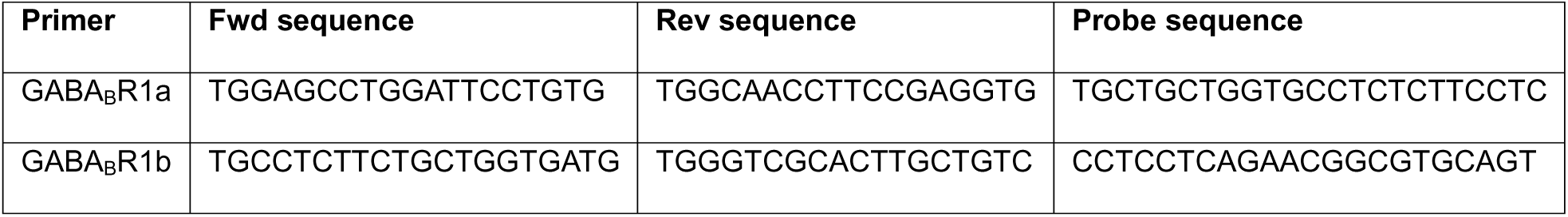
Designed isoform-specific primer sequences

## Results

### Oligodendrocytes show the highest levels of APP RNA

APP is broadly expressed throughout the brain, yet its relative expression across different cell types and the implications of this expression remain underexplored. We examined the relative RNA expression of APP (gene name *App*) in a publicly available RNAseq dataset generated by Zhang et al.^30^ which isolated oligodendrocytes, microglia, endothelial cells, astrocytes, and neurons using a variety of cell-sorting and isolation methods. *App* was most highly expressed in oligodendrocytes followed by endothelial cells and neurons (Figure 1A). While this dataset included all cell types of interest, samples were prepared using different cell isolation methods and mouse genotypes. Therefore, we performed magnetic-activated cell sorting (MACS) to sequentially isolate oligodendrocytes, microglia, endothelial cells, astrocytes, and neurons from the same mouse brain^29^ and probed for RNA expression of APP using RT-qPCR. Primers were validated using APP KO mice (Supplemental figure 1). The purity and viability of the sorted cell fractions were validated using flow cytometry and RT-qPCR, as reported in Houmam et al., 2025^29^. Our results corroborated those shown in the publicly available RNAseq dataset^30^ . *App* was strongly expressed in oligodendrocytes in both males (1B) and females (1C) followed by endothelial cells and neurons (Figure 1B and C). These results suggest that APP signaling may play a major role in oligodendrocytes.

**Figure 1:**
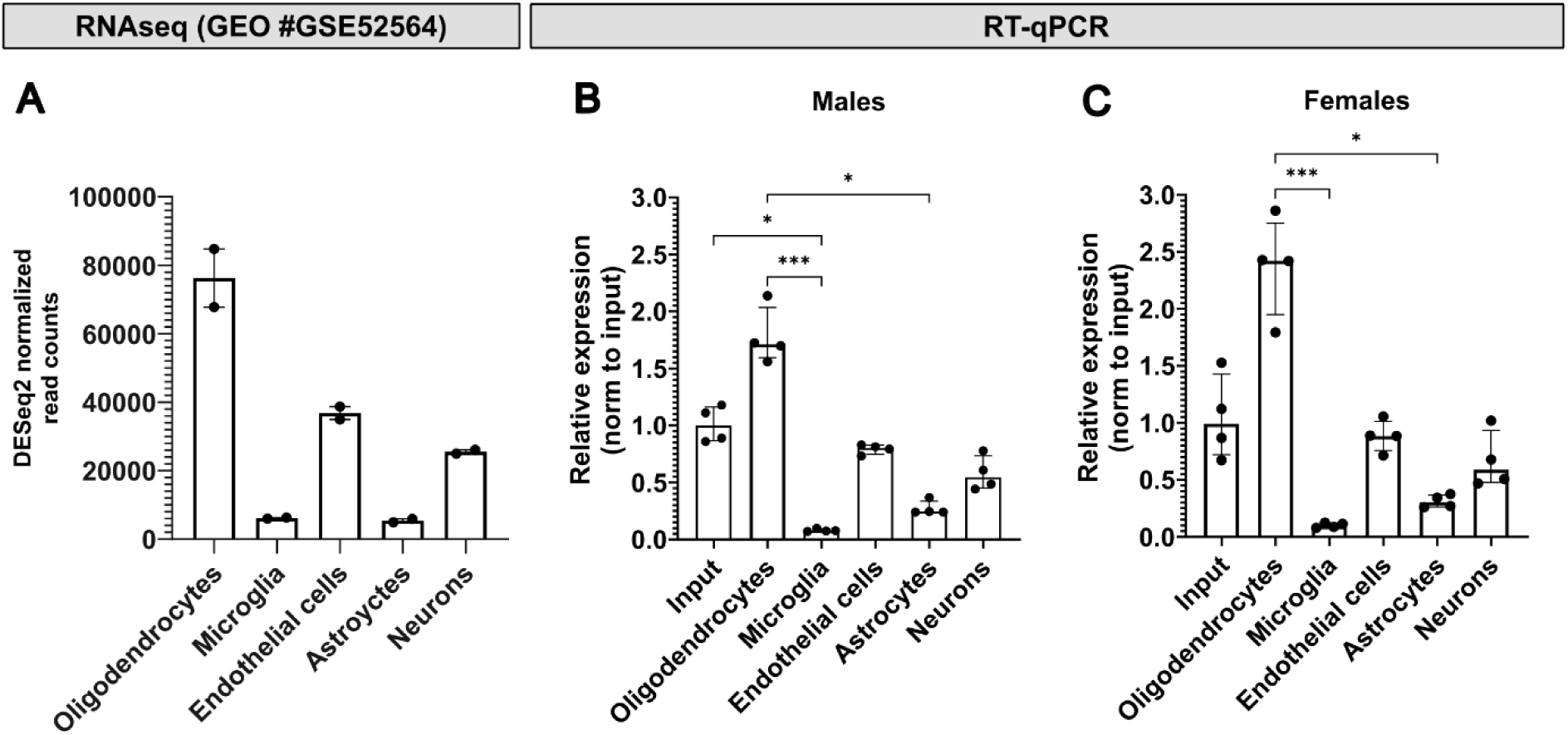
APP RNA expression levels in oligodendrocytes, microglia, endothelial cells, astrocytes, and neurons **(A)** DESeq2 normalized read counts for APP in oligodendrocytes, microglia, endothelial cells, astrocytes, and neurons from GEO #GSE52564 (Kruskal-Wallis P=0.0011; Dunn’s multiple comparisons: all ns due to small sample size, n=2 per group). **(B)** Relative APP RNA expression measured by RT-qPCR in sorted cells from male mice (Kruskal-Wallis P=0.0005; Dunn’s multiple comparisons: * = p≤0.05, *** = p≤0.001; n=4 per group). **(C)** Relative APP expression in sorted cells from female mice (Kruskal-Wallis P=0.0010; Dunn’s multiple comparisons as in B; n=4 per group). Data shown as median ± IQR.

### Oligodendrocytes show the highest GABA_B_R1a RNA levels

GABA_B_R is an obligate heterodimer composed of subunit GABA_B_R1 which contains the binding site for GABA, its endogenous agonist, and subunit GABA_B_R2 which is linked intracellularly to G-proteins. Alternative promoter usage can result in two predominant isoforms of the GABA_B_R1 subunit referred to as GABA_B_R1a and GABA_B_R1b. The GABA_B_R1a isoform contains two N-terminal sushi repeats not present in subunit GABA_B_R1b (Figure 2A and B). Sushi domain 1 is required for APP-GABA_B_R binding and therefore this interaction requires the GABA_B_R1a isoform to function^12,13^. While both GABA_B_R1 and GABA_B_R2 subunits have been shown to be expressed in neurons and glial cells, whether these cells express both the GABA_B_R1a and 1b isoform equally is still unclear. Therefore, we sought to determine the isoform-specific expression patterns of GABA_B_R across cell types.

**Figure 2:**
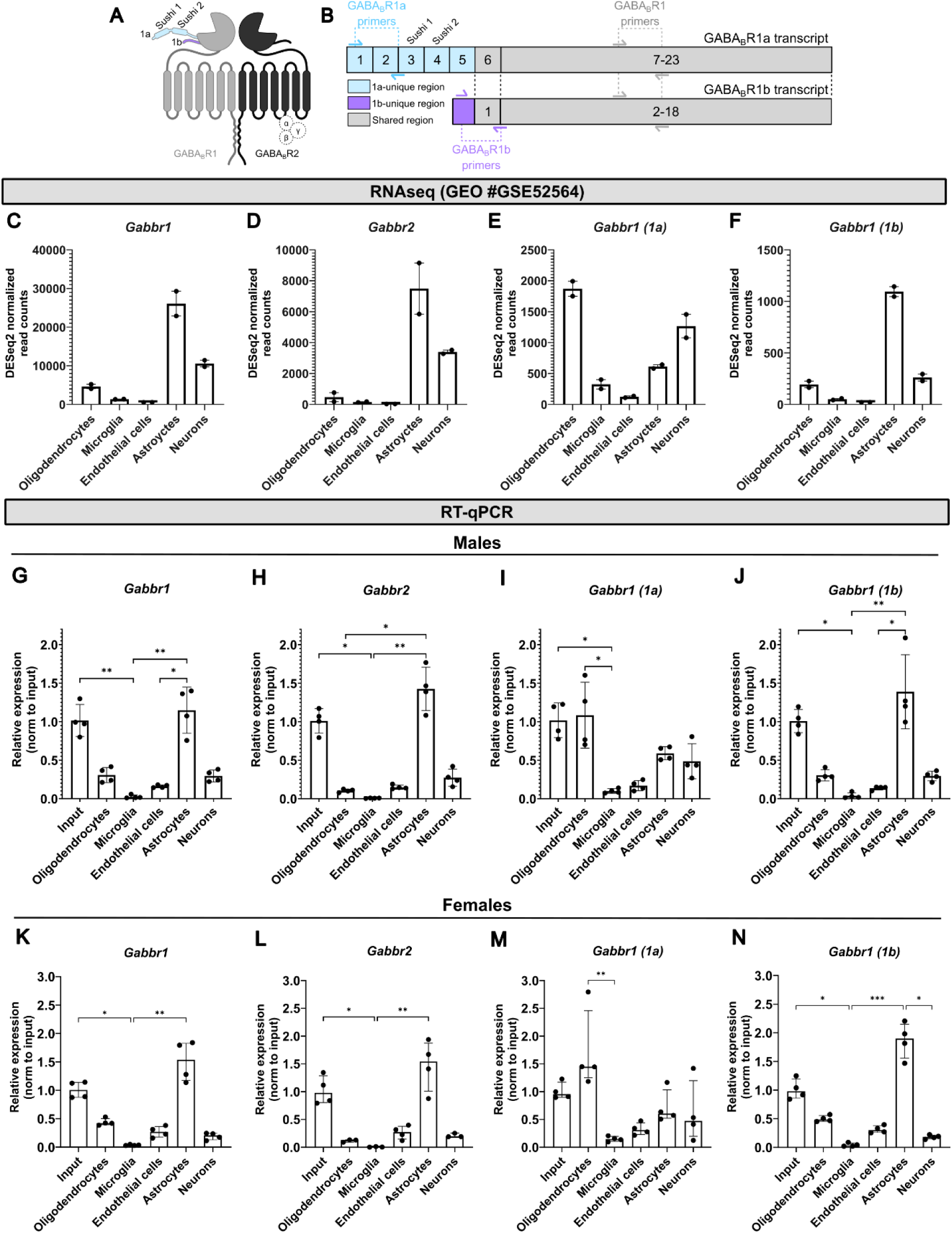
GABA_B_R isoform-specific RNA expression levels in oligodendrocytes, microglia, endothelial cells, astrocytes, and neurons **(A)** Schematic of GABA_B_R subunits and isoforms. The GABA_B_R1a isoform contains sushi domains (blue) absent in GABA_B_R1b (purple). **(B)** Schematic of GABA_B_R1a and GABA_B_R1b transcripts: GABA_B_R1a contains five unique exons (blue), while GABA_B_R1b contains only a short unique sequence at the start of exon 1 (purple). **(C–F)** DESeq2 normalized read counts from GEO #GSE52564 for GABA_B_R1 (C, P=0.0011), GABA_B_R2 (D, P=0.0053), GABA_B_R1a (E, P=0.0011), and GABA_B_R1b (F, P=0.0011) across oligodendrocytes, microglia, endothelial cells, astrocytes, and neurons (Kruskal-Wallis. Dunn’s multiple comparisons all ns due to small sample size, n=2). **(G–N)** Relative RNA expression measured by RT-qPCR in sorted oligodendrocytes, microglia, endothelial cells, astrocytes, and neurons from male (G–J) and female (K–N) WT mice. Statistical significance was determined by Kruskal–Wallis test (Males: Gabbr1, P=0.0007; Gabbr2, P=0.0005; 1a, P=0.0013; 1b, P=0.0007; Females: Gabbr1, P=0.0006; Gabbr2, P=0.0028; 1a, P=0.0031; 1b, P=0.0004), followed by Dunn’s multiple comparisons (* = p ≤ 0.05; ** = p ≤ 0.01; *** = p ≤ 0.001). Data shown as median ± IQR; n=4 per group.

Using the publicly available dataset generated by Zhang et al.^30^, we probed for the RNA expression of total GABA_B_R1 and GABA_B_R2 and found that both subunit RNA levels are highest in astrocytes followed by neurons and oligodendrocytes (Figure 2C and D). To be able to distinguish read counts for the GABA_B_R1a and GABA_B_R1b isoforms of the GABA_B_R1 subunit, we developed an isoform-aware workflow by aligning read counts generated by the study to a manually edited genome file where sequences unique to each isoform transcript were labeled *Gabbr1 (1a)* and *Gabbr1 (1b)* (Figure 2B).

We found that total GABA_B_R1 RNA was highest in astrocytes. However, RNA levels of the GABA_B_R1a subunit were highest in oligodendrocytes, followed by neurons, astrocytes, and microglia (Figure 2E). On the other hand, GABA_B_R1b RNA was highest in astrocytes, followed by neurons and astrocytes (Figure 2F).

The unique isoform-specific regions of GABA_B_R1a and GABA_B_R1b are short, GC rich, and located at the 5’ end of the transcripts (Figure 2B). Since these features result in low read counts in RNAseq^32,33^, which is typically 3’- biased and inefficient at detecting, short, GC-rich sequences, we then employed a more targeted approach. We performed RT-qPCR with commercially available primers to assess expression of total GABA_B_R1 and GABA_B_R2 and custom-designed primers to assess isoform-specific GABA_B_R1a and GABA_B_R1b levels (Figure 2B). We validated our custom primers using a GABA_B_R1a-specific KO mouse model. The GABA_B_R1a primer was specific to the R1a isoform, showing no amplification in GABA_B_R1a KO samples (Supplemental figure 2A). As expected, primers targeting the 1b isoform showed no change in expression between the GABA_B_R1a KO and WT. (Supplemental figure 2B). Total GABA_B_R1 RNA levels were highest in astrocytes followed by oligodendrocytes and neurons in both male (Figure 2G) and female (Figure 2K) mice. GABA_B_R2 RNA levels were highest in astrocytes followed by neurons in both males (Figure 2H) and females (Figure 2L). Isoform-specific RT-qPCR showed that RNA levels of the GABA_B_R1a subunit isoform were highest in oligodendrocytes followed by astrocytes and neurons in males (Figure 2I) and females (Figure 2M), and the GABA_B_R1b was highest in astrocytes followed by oligodendrocytes and neurons in male (Figure 2J) and female mice (Figure 2N). These findings suggest a cell-type bias, with GABA_B_R1a-mediated signaling prominent in oligodendrocytes and GABA_B_R1b-mediated signaling predominating in astrocytes. Moreover, our findings highlights oligodendrocytes as a cell type where both APP and GABA_B_R1a RNA are most highly expressed in the brain.

### APP KO does not affect GABA_B_R1a transcript levels

Lastly, we tested whether APP KO may affect transcript levels of GABA_B_R1 due to a compensatory mechanism. Using APP KO mice (Supplemental figure 1) we assessed overall levels of GABA_B_R1 as well as GABA_B_R1a and GABA_B_R1b. APP KO did not affect levels of GABA_B_R1 and GABA_B_R1a in any of the cell types (Figure 3B, C). Intriguingly, GABA_B_R1b was increased in APP KO astrocytes (Figure 3D). This may be due to an astrocyte-specific compensation to the lack of GABA_B_R1a signaling, and may hint at GABA_B_R function being particularly important in astrocytes. We also tested whether GABA_B_R1a KO may alter RNA levels of APP. APP RNA levels were unchanged in all five cell types in GABA_B_R1a KO mice (Figure 3A).

**Figure 3:**
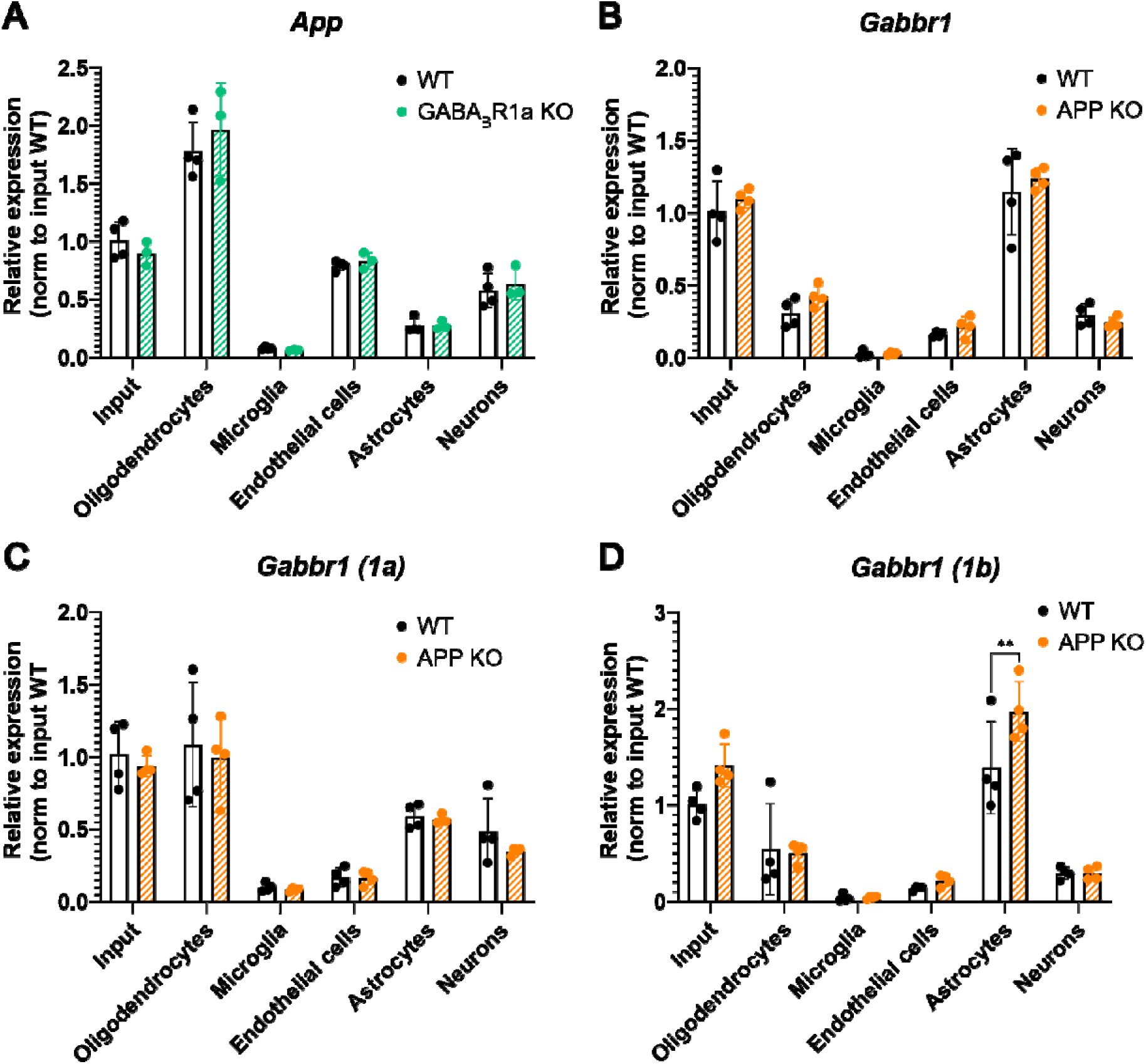
Cell-type specific expression levels of APP and GABA_B_R1 in knockout mouse models Relative RNA expression measured by RT-qPCR in sorted oligodendrocytes, microglia, endothelial cells, astrocytes, and neurons. **(A)** APP expression in WT (black) and GABABR1a KO (green) mice. **(B–D)** Gabbr1 (B), Gabbr1 (1a) (C), and Gabbr1 (1b) (D) expression in WT (black) and APP KO (orange) mice. Two-way ANOVA with factors genotype and cell type was performed, followed by Sidak’s multiple comparisons between genotypes within each cell type (** = p ≤ 0.01). Data are presented as mean ± SD; n=4 per group.

## Discussion

While APP expression has been documented in numerous brain cell types, the extent and functional implications of its expression across different non-neuronal cell types is still underexplored. Our results confirm that APP is expressed not only in neurons but also in oligodendrocytes, microglia, endothelial cells, and astrocytes. We show that APP is most highly expressed in oligodendrocytes, a cell type in which it remains largely understudied. APP and its cleavage fragments appear to modulate oligodendrocyte-led processes such as myelination and re-myelination of axons^8,34^. Promoting sAPPα production can restore damaged myelin and protect against further demyelination by increasing myelin basic protein and mature oligodendrocyte levels in the cerebellum^35^. Oligodendrocytes may play a critical role in AD pathology, as myelin disruption has been identified as a feature of the disease^36^. Moreover, oligodendrocytes have recently been emphasized as a significant source of Aβ peptides further underlining their potential involvement in AD^37–39^. We also show that endothelial cells express the second-highest levels of APP among brain cell types. This aligns with earlier work showing that APP is expressed in endothelial cells where it contributes to vascular integrity and nitric oxide signaling^11,40,41^.

Similar to APP, GABA_B_R is well characterized in neurons but largely understudied in glial cells. While GABA_B_R is known for its primary role in modulating synaptic inhibition in neurons, its expression is not limited to neurons^22,25,26^. We validate this by showing that overall expression levels of GABA_B_R1 and GABA_B_R2 in the brain are highest in astrocytes. However, when we assessed isoform-specific expression of the GABA_B_R1 subunit, we found that RNA levels of the GABA_B_R1a subunit are highest in oligodendrocytes while the GABA_B_R1b subunit is highest in astrocytes. This suggests that processes mediated by GABA_B_R1a specifically such as APP-mediated GABA_B_R signaling may be important in oligodendrocytes where both APP and GABA_B_R1a are expressed at high levels. GABA_B_R has been shown to be expressed in oligodendrocyte lineage cells and are regulated during OPC differentiation to myelinating oligodendrocytes^23,42^. Specifically, GABA_B_R1 activation using Baclofen is reported to promote remyelination in the brain^24^. It remains unclear whether these GABA_B_R-mediated oligodendrocyte functions are through one or both isoforms, given that Baclofen is not an isoform-specific agonist. Our data hints at a GABA_B_R1a-dominant oligodendrocyte signaling, however further experimentation is needed to validate this finding mechanistically. An isoform specific interactor of GABA_B_R1 such as sAPP, therefore, provides a highly useful tool to tease apart R1a-specific effects of GABA_B_R activation^43^.

Our isoform-specific results show that the GABA_B_R1b subunit isoform is enriched in astrocytes. In addition, astrocytes were the only cell type to upregulate expression of GABA_B_R1b in the APP KO mouse model. GABA_B_Rs have been shown to be present on astrocytes where they can modulate their inflammatory response to stimuli such as lipopolysaccharide^26^. Stimulation of GABA_B_Rs in astrocytes regulates adenylyl cyclase activity^44^ and mediates calcium signaling in astrocytes which supports interneuron activity^45^. Our data suggests that these GABA_B_R-mediated astrocytic functions may be through the GABA_B_R1b isoform, but further mechanistic work using a combination of baclofen and sAPP stimulation is needed to elucidate this.

Lastly, our results show that the expression pattern of GABA_B_R2 RNA follows that of overall GABA_B_R1 closely in the five cell types. However, when assessing isoform-specific expression of GABA_B_R1, we observed that GABA_B_R2 relative levels match those of GABA_B_R1b but not GABA_B_R1a. Specifically, oligodendrocytes seem to express significantly high levels of GABA_B_R1a but are among the lowest expressing cell types for GABA_B_R2. Current consensus in the field indicates that GABA_B_Rs are obligate heterodimers of GABA_B_R1 and GABA_B_R2 subunits, with only a few reports challenging this model^20^. When expressed alone, GABA_B_R1 has been shown to illicit smaller responses suggesting it could be at least partially functional alone or in combination with an unknown partner^20,46^. In oligodendrocytes, GABA_B_R1 levels may be regulated by differentiation stage where the ratio of GABA_B_R1 to GABA_B_R2 is thought to be changed^23^. Cell-type-specific studies of GABA_B_R signaling could reveal whether R1/R2 heterodimerization is consistent across all brain cells and whether both subunits are required for sAPP-GABA_B_R signaling.

To summarize, we report oligodendrocytes as a cell type where both APP and GABA_B_R1a RNA are highly expressed using both analysis of a publicly available dataset^30^ and RT-qPCR of cell-sorted mouse brains. Our findings highlight oligodendrocytes as a novel potential cellular location for the APP-GABA_B_R interaction to be investigated. Future studies can build on this work by mapping GABA_B_R cell-specific isoform expression across brain regions and developmental stages, confirming protein-level expression, and testing the functional impact of APP-GABA_B_R signaling in oligodendrocytes on key processes such as myelination of neurons.

## Supporting information

Graphical Abstract

## Author contributions

S.H., D.S., M.S., C.G.M., and H.C.R. conceived the study. S.H. and D.S. performed RT-qPCR experiments and data analysis. S.H., M.S, G.B.H., N.P.P., and D.R.S. designed and performed isoform-specific RNAseq analysis of publicly available dataset. C.S. and Y.M.T. provided technical assistance. S.H. and H.C.R wrote the manuscript. D.S., D.R.S., and G.B.H. edited the manuscript.

## Data availability statement

The data that support the findings of this study are available from the corresponding author upon reasonable request. The publicly available RNAseq dataset analysed in this paper is accessible under the following GEO accession number: <u>GSE52564</u>.

## Ethics approval statement

All animal procedures described in this protocol were approved by the Institutional Animal Care and Use Committee of the Oklahoma Medical Research Foundation and performed in accordance with the National Institutes of Health (NIH) Guide for the Care and Use of Laboratory Animals

## Acknowledgements

This work was supported by the National Institutes of Health (R35GM142726, R21AG085486, P20GM125528 to H.C.R. and T32AG052363 to S.H) and the Presbyterian Health Foundation (Pilot Research Funding to H.C.R.). Data processing and analysis were supported by the OMRF Center for Biomedical Data Sciences. We thank the OMRF flow cytometry core and members of the Rice, Freeman, and Ocañas, and Miller labs for helpful discussions. We thank the Freeman lab for providing access to the MACSOctodissociator and Quantstudio5 instruments. We thank the Lewis lab for providing access to the QuadroMACS stand The TOCI diagrams were created in BioRender. Houmam, S. (2026) https://BioRender.com/zugxqwy.

