## Supplemental Figures for "Oligodendrocytes show enriched expression of amyloid precursor protein and GABA B receptor isoform 1a"

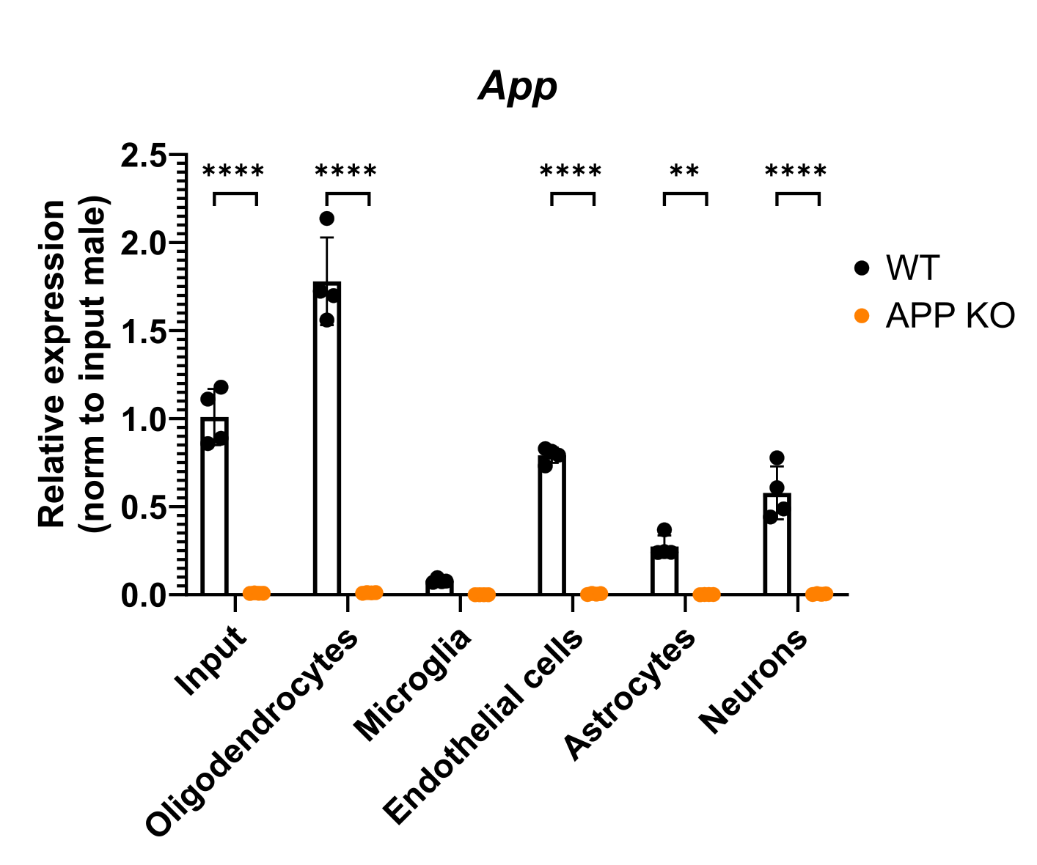

**Supplemental figure 1: Validation of RT-qPCR primer for APP**

Relative APP RNA expression measured by RT-qPCR in sorted oligodendrocytes, microglia, endothelial cells, astrocytes, and neurons from WT (black) and APP KO (orange) mice. (Two-way ANOVA was performed with factors genotype and cell type, followed by Sidak’s multiple comparisons between genotypes within each cell type (** = p≤0.01, ** = p≤0.0001). n=4 per group). Data shown as mean ± SD.

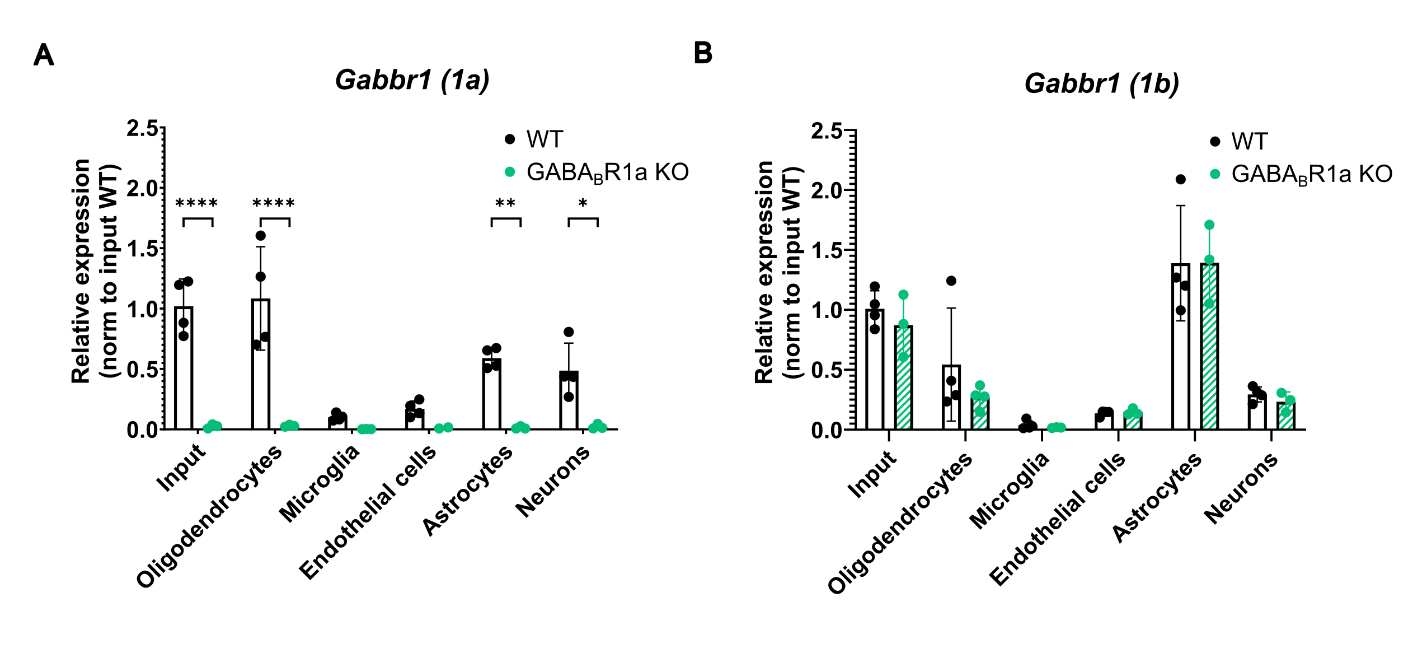

**Supplemental figure 2: Validation of isoform-specific RT-qPCR primers for GABA_B_R1**

Relative RNA expression of **(A)** Gabbr1 (1a) and **(B)** Gabbr1 (1b) measured by RT-qPCR in sorted oligodendrocytes, microglia, endothelial cells, astrocytes, and neurons from WT (black) and GABABR1a KO (green) mice. Two-way ANOVA with factors genotype and cell type was performed, followed by Sidak’s multiple comparisons between genotypes within each cell type (* = p ≤ 0.05; ** = p ≤ 0.01; **** = p ≤ 0.0001). Data are presented as mean ± SD; n=4 per group.
