## Supplementary figures and images for "Oligodendrocytes show enriched expression of amyloid precursor protein and GABA B receptor isoform 1a"

### Graphical Abstract

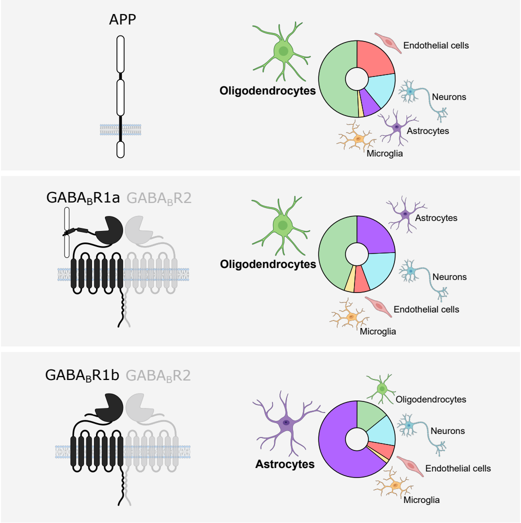
